# Mapping load-bearing in the mammalian spindle reveals local kinetochore-fiber anchorage that provides mechanical isolation and redundancy

**DOI:** 10.1101/103572

**Authors:** Mary Williard Elting, Manu Prakash, Dylan B. Udy, Sophie Dumont

## Abstract

Active forces generated at kinetochores move chromosomes, and the dynamic spindle must robustly anchor kinetochore-fibers (k-fibers) to bear this load. We know that the mammalian spindle body can bear the load of chromosome movement far from poles, but do not know where and how – physically and molecularly – this load is distributed across the spindle. In part, this is because perturbing and reading out spindle mechanics in live cells is difficult. Yet, answering this question is key to understanding how the spindle generates and responds to force, and performs its diverse mechanical functions. Here, we map load-bearing across the mammalian spindle in space-time, and dissect local anchorage mechanics and mechanism. To do so, we laser ablate single k-fibers at different spindle locations, and in different molecular backgrounds, and quantify at high time resolution the immediate relaxation of chromosomes, k-fibers, and microtubule speckles. We find that load redistribution is locally confined in all directions: along the first 3-4 μm from kinetochores, scaling with k-fiber length, and laterally within ~2 μm of k-fiber sides, without neighboring k-fibers sharing load. A phenomenological model constrains the mechanistic underpinnings of these data: it suggests that dense, transient crosslinks to the spindle along k-fibers bear the load of chromosome movement, but that these connections do not limit the timescale of spindle reorganization. The microtubule crosslinker NuMA is needed for the local load-bearing observed, while Eg5 and PRC1 are not, suggesting specialization in mechanical function and a novel function for NuMA throughout the spindle body. Together, the data and model suggest that widespread NuMA-mediated crosslinks locally bear load, providing mechanical isolation and redundancy while allowing spindle fluidity. These features are well-suited to support robust chromosome segregation.

## Introduction

When mammalian cells divide, depolymerizing microtubules generate forces at kinetochores to move chromosomes and ultimately segregate them. Microtubule bundles called kinetochore-fibers (k-fibers) attach to kinetochores; thus equal and opposite forces are exerted on kinetochores by k-fibers and on k-fibers by kinetochores, in accord with Newton’s Third Law (Fig. 1a). However, for chromosomes – rather than k-fibers – to move, k-fibers must anchor to the spindle to bear the load of chromosome movement [1]. How the mammalian spindle robustly anchors k-fibers and segregates chromosomes despite its dynamic and flexible nature remains unclear. Yet, answering this question is key to understanding how the spindle generates and responds to force, and thus to determining how the cell moves chromosomes, and regulates their attachment and segregation.

**Figure 1.**
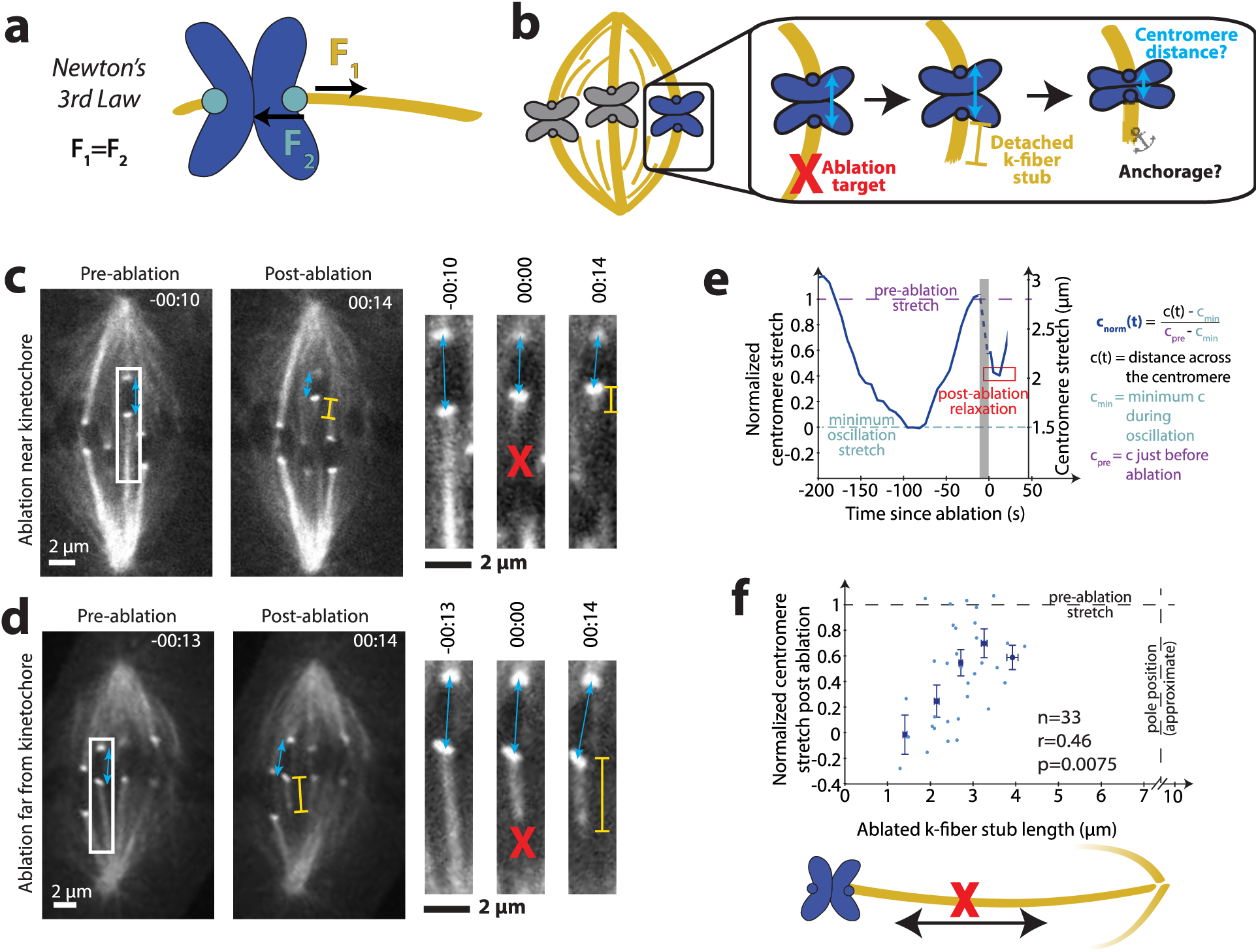
The load of chromosome movement is borne proportionally along the first 3-4 μm of the k-fiber. See also Videos 1 and 2. **(a)** Equal and opposite forces pull on kinetochores (cyan) and k-fibers (yellow). **(b)** Cartoon of ablation assay to map where the spindle bears load along the k-fiber. Centromere stretch that remains after ablation indicates residual force on the k-fiber stub. **(c)** Live confocal imaging of Ptk2 cell expressing EYFP-Cdc20 and EGFP-α-tubulin. Time since ablation, min:sec. Representative example showing that when we ablate (red X) near kinetochores, the centromere (blue arrow) relaxes (inset), indicating decreased load-bearing ability by the short k-fiber stub (yellow bar). **(d)** Imaging as in (c), representative example showing that after ablation far from kinetochores, the centromere remains stretched, indicating preserved load-bearing ability by the long k-fiber stub. **(e)** Representative plot of centromere stretch over time before and after ablation, normalized as defined on a per-trace basis (see Exp. Proc.). **(f)** Normalized centromere stretch post-ablation vs. k-fiber stub length of individual measurements (light blue dots) and binned data (darker, larger dots; error bars s.e.m.).

In one model, k-fibers are anchored at spindle poles, and their microtubules are tensed all along their lengths [2–4]. However, k-fibers can bear load and support chromosome movement without being directly connected to the pole or other k-fibers in both mammalian [5, 6] and insect cells [7]. These observations support a model where k-fibers are anchored along their lengths: in this model, different sections of the k-fibers can be under different forces [5, 8], and “traction force” may [9] or may not [10] be proportional to k-fiber length.

The physical and molecular bases of k-fiber anchorage in the spindle body are poorly understood. The local physical environment (i.e., viscoelastic properties) around a k-fiber dictates how kinetochore forces are distributed in space and time across the spindle; yet, while directly probing local mechanics has become possible in spindles that can be reconstituted in extract [11, 12], such perturbation remains challenging in mammalian cells. Although we do not know what structure k-fibers anchor to, spindle microtubules connecting to k-fibers [13] are a clear candidate. These connecting microtubules could, for example, include interpolar microtubules [8, 14, 15], “bridging fiber” microtubules [5], and neighboring k-fibers connecting through minus-ends [16, 17] – or other mechanisms could be responsible [18]. A further challenge is that if the spindle laterally bears load, it must do so using an anisotropic spindle material that is weaker along its lateral axis [8, 11, 13, 19–21]. The first identification herein of a molecule whose absence changes the spindle’s load-bearing map is an important step towards uncovering the mechanisms that bear load – and understanding how they support the mechanical integrity and dynamics essential to accurate chromosome segregation.

Here, we combine targeted k-fiber laser ablation and high resolution imaging to build a map of where the mammalian spindle bears the load of chromosome movement and to probe the mechanics and molecular basis of the underlying anchorage. We find that effective anchorage is proportional to length in the first few microns of a k-fiber, but does not require the entire length from kinetochore to pole, and that neighboring k-fibers do not share the load. The data support a model where spindle connections linearly distributed along k-fibers bear load, but where the viscosity of these connections does not limit the timescale of spindle reorganization. We find that the microtubule crosslinker NuMA is essential for locally bearing the load of chromosome forces, while other molecules linking microtubules, Eg5 and PRC1, are not. This dynamic and local nature of load-bearing is well-suited for making chromosome movement robust to spindle reorganization and for spatially confining the impact of structural spindle defects.

## Results & Discussion

### The load of chromosome movement is borne proportionally along the first 3-4 μm of the k-fiber

To map where along their lengths k-fibers bear the load of chromosome movement in mammalian spindles, we used laser ablation to sever k-fibers at varying distances from the kinetochore (Fig. 1b). We visualized both microtubules and kinetochores (EGFP-α-tubulin and EYFP-Cdc20, respectively) in Ptk2 cells, which have k-fibers whose microtubules largely extend from kinetochore to pole [22], and have few chromosomes, allowing us to effectively target individual k-fibers. We began by measuring the response of the chromosome since, at metaphase, forces at kinetochores stretch centromeres and are opposed by k-fiber anchorage in the spindle. Thus, high residual centromere stretch after ablation reflects strong anchorage on the remaining k-fiber stub, whereas low residual stretch indicates weak anchorage. We ablated k-fibers at times of maximal centromere stretch to maximize the dynamic range of the assay, and examined the new steady state centromere distances with ablated, detached k-fiber stubs. This new, brief steady state occurred before the ablated k-fiber stubs were pulled poleward by dynein [16, 17].

We systematically ablated k-fibers from 1 to 4 μm from kinetochores and mapped how centromere relaxation changed with ablation position. The centromere relaxed when ablation was near the kinetochore (Fig. 1c, Video 1), and remained stretched when ablation was farther from the kinetochore, but also far from poles (Fig. 1d, Video 2). This observation is in qualitative agreement with previous work in mammalian cells [5, 6], and indicates that long k-fiber stubs remained anchored following ablation, without a direct connection to the pole or other k-fibers. However, here we see a considerable drop in centromere stretch for stubs as long as 2-3 μm, suggesting that load is borne gradually over a few microns. The normalized centromere distance (Fig. 1e) to which each chromosome relaxed correlated with the distance from the kinetochore to the ablation site (r=0.46, p=0.0075, n=33) (Fig. 1f). Ablating a single k-fiber, maximizing the dynamic range of our assay, and normalizing for centromere variability are likely key to revealing how load is distributed along k-fiber lengths. The apparent linear relationship, which occurs for stubs between ~1 and ~3 μm in length, suggests that load-bearing along the k-fiber stub scales with its length. However, this relationship plateaus: for stubs at least 3-4 μm in length, we observed little relaxation following ablation (note that it is difficult to ablate more than 4 μm away and still target single k-fibers, since they come closer together near poles). Further, k-fiber stubs of <~1.5 μm were not able to generate sufficient load to stretch the centromere beyond its minimal metaphase distance. Together, the data are consistent with a constant number of molecular-scale force generators per unit length which bear load locally, in the first 3-4 μm near the kinetochore, and inconsistent with a single anchor point at a fixed distance from the kinetochore.

### The load of chromosome movement is not distributed laterally to neighboring k-fibers and is confined to non-k-fibers less than ~2 μm away

To probe how the load of chromosome movement is distributed in the spindle body, we mapped load sharing along the lateral spindle axis. We reasoned that if lateral anchorage distributed significant load to neighboring k-fibers (as has been previously proposed [14, 23]), we should see less effective anchorage of k-fibers at the spindle edge, which have fewer neighbors, compared to those in the spindle center, which have more neighbors. We separated the previous data set (Fig. 1f) into “edge” and “center” k-fibers and could not detect a significant difference in centromere relaxation after ablation in these two populations (Fig. 2a; n=14 and 19, respectively). This similarity in behavior suggests that lateral load-bearing also occurs locally.

**Figure 2.**
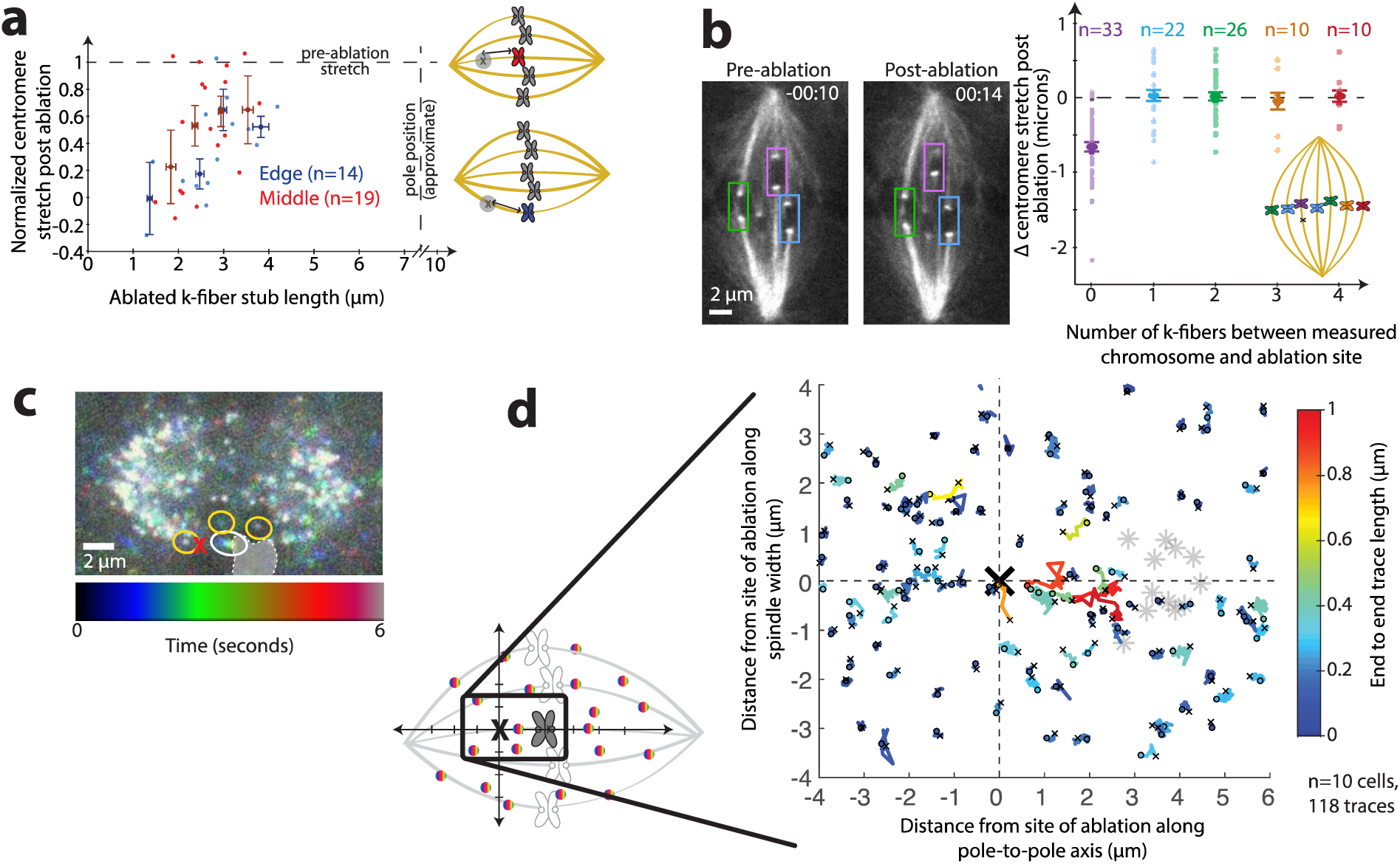
The load of kinetochore forces is borne laterally within ~2 μm of k-fibers. See also Video 3. **(a)** Normalized centromere stretch vs. k-fiber stub length of individual measurements (light dots) and binned data (darker, larger dots; error bars s.e.m.) for chromosomes on the edge (blue) and in the center (red) of the spindle with same source data as Fig. 1f. **(b)** Left, representative live imaged Ptk2 cell (same cell as in Fig. 1c) diagramming neighboring chromosomes. Time since ablation, min:sec. Right, change in centromere stretch following ablation for ablated k-fiber (purple) and increasingly distant neighboring k-fibers across the spindle (blue, green, orange, and red, as in cartoon). Individual measurements, lighter dots. Binned data, with s.e.m. error bars, larger dots. **(c)** Live confocal movie of Ptk2 cell expressing SunTag-rigor kinesin speckles at a fast time scale (~300 ms), projected into a single plane colored with time. A speckle on the ablated k-fiber (white circle) appears as a rainbow because it relaxes a significant distance during the movie, whereas nearby speckles (e.g., yellow circles) do not move, and thus appear white. Chromosome position at first frame indicated by shaded region. **(d)** Individual speckle tracks (colored by their end-to-end distance) and chromosomes whose k-fibers were targeted (gray stars) are shown superimposed and aligned, with orientation as in cartoon (see Supp. Exp. Proc.). The majority of long speckle traces (warm colors) representing significant relaxation are between the site of ablation and the chromosome, whereas speckles >~2 microns laterally away from the site of ablation typically only move short distances (cool colors). Small ‘o’ indicates beginning of each trace and small ‘x’ indicates end.

To further test whether neighboring k-fibers shared their load, we ablated a single k-fiber and examined the responses of centromeres of neighboring chromosomes following ablation. If significant force were distributed between k-fibers, cutting one k-fiber would affect the anchorage of neighboring k-fibers, causing relaxation of their centromeres. From distant k-fibers to nearest neighbor k-fibers, we could not detect a significant decrease in centromere stretch following ablation (Fig. 2b, n=68). This observation, too, indicates that that the load of k-fiber anchorage is distributed over a shorter distance than the distance between neighboring k-fibers (~2 μm), and thus, that spindle structures located in between these k-fibers must bear the load of chromosome movement. Consistently, prior measurements found only small correlations between the movement of neighboring chromosomes [18], which may result from closer neighbor k-fibers in human cells, or from chromosome/chromosome (rather than k-fiber/k-fiber) interactions. Thus, neighboring k-fibers act as mechanically decoupled bodies in the spindle.

To directly map how microtubules across the whole spindle mechanically respond to local changes in architecture and force, we imaged microtubule speckles at high time resolution (~300 ms) after k-fiber laser ablation. We sparsely labeled microtubules by expressing a Sun-tagged truncated kinesin-1 mutant that rigor binds to microtubules [24]. This bright, single molecule imaging method allowed us to visualize the movement of microtubules with respect to each other. After ablation, speckles apparently located on the ablated k-fiber moved away from the ablation site at similar speeds to whole k-fibers. We only observed considerable relaxation for speckles within ~3 μm of the site of ablation. While most of these speckles were presumably localized along ablated k-fibers, we also observed speckle relaxation within ~2 μm from the site of ablation along the lateral axis, apparently next to the ablated k-fibers on non-k-fiber microtubules. Speckles farther away had no detectable movement (Fig. 2c, d, Video 3, n=118 speckles in 10 cells). Thus, a k-fiber is not only mechanically isolated from neighboring k-fibers, but also from non-k-fiber microtubules more than ~2 μm away. Relaxation of nearby speckles adjacent to the ablated k-fiber is consistent with these k-fibers anchoring to neighboring non-k-fiber microtubules. In contrast, the fixed appearance of speckles farther away suggests that spindle deformations do not propagate far across the spindle and do not undermine its overall microtubule architecture. Together, the data suggest that this mechanical isolation is, in part, accomplished by a spindle structure that propagates stresses and strains predominately along a pole-to-pole, rather than a lateral, axis, and does so locally.

### K-fiber post-ablation relaxation dynamics and steady state support a viscoelastic model of the connection of the k-fiber to the spindle

To probe the mechanics of the connections between k-fibers and the spindle – the local spindle environment a k-fiber experiences – we asked how relaxation dynamics following ablation varied with k-fiber stub length. We imaged at faster time resolution (~300 ms) than in Fig. 1, and used Ptk2 GFP-tubulin cells as they provide good imaging contrast during fast imaging despite photobleaching. We tracked the kinetochore ends of the k-fiber stubs over time (Fig. 3a, Video 4). Most of the relaxation responses we observed fit well to single exponentials (timescales *τ*, Fig. 3b). An exponential form is consistent with viscoelastic relaxation, and smooth relaxation is consistent with many weak interactions, rather than a few strong ones, anchoring k-fibers. We did not observe a significant correlation between the lengths of the k-fiber stubs and the timescales of centromere relaxation (Fig. 3c) (r=0.17, p=0.6, n=12); similarly, previous results with only short stubs also found that relaxation of the kinetochore itself did not depend on stub length [25].

**Figure 3.**
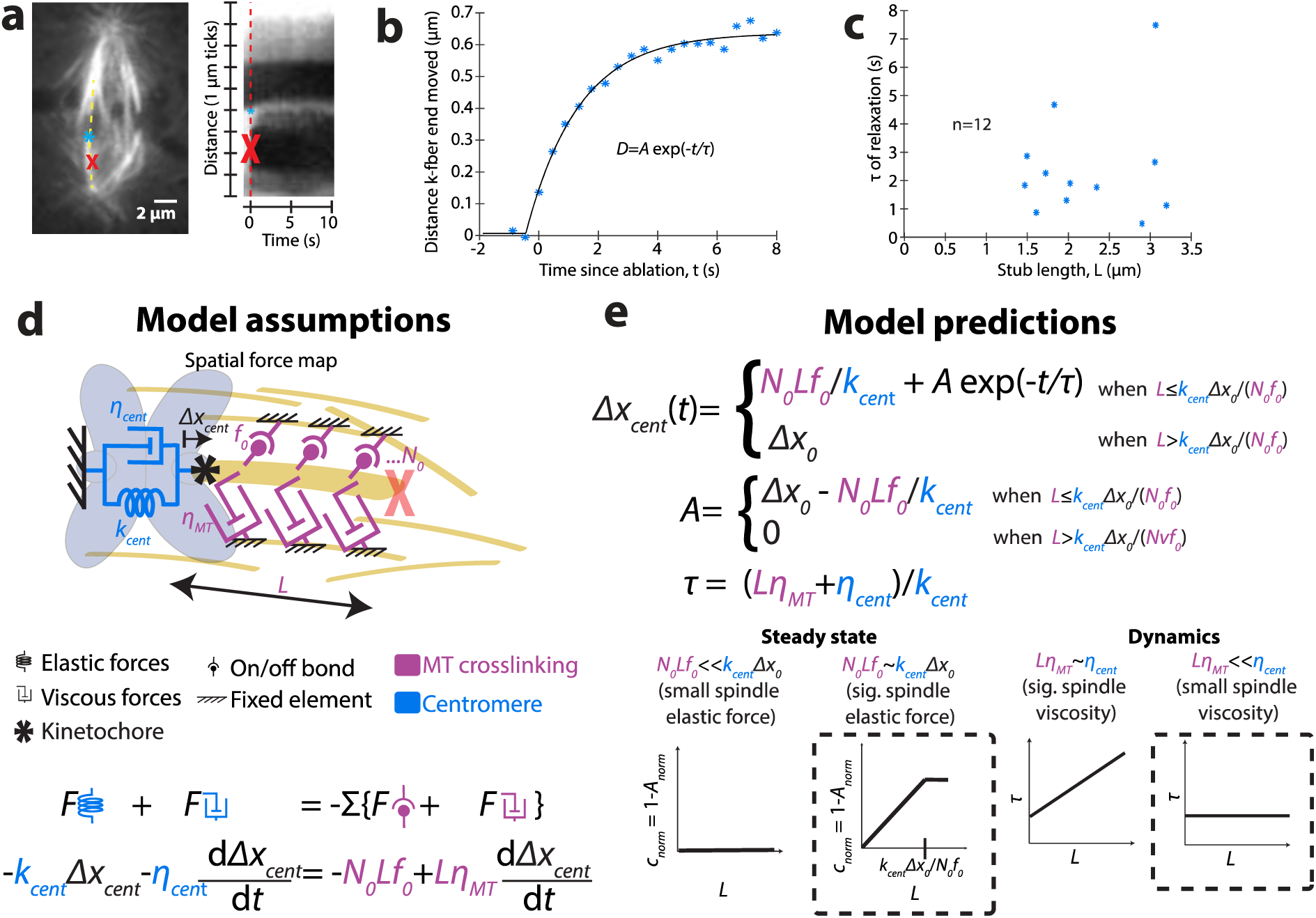
K-fiber post-ablation relaxation dynamics and steady state support a viscoelastic model of the connection of the k-fiber to the spindle. See also Video 4. **(a)** Live confocal image immediately before ablation (at red X) of Ptk2 cell expressing EGFP-α-tubulin. Kymograph (right) along a k-fiber (yellow dotted line) of imaging at fast time resolution (~300 ms) shows the relaxation process of the tracked k-fiber end (blue *). **(b)** Tracking of relaxing k-fiber stub shown in (a) over time, with fit to an exponential relaxation. **(c)** Time scales of relaxation (from exponential fits) as a function of stub length (r=0.17, p=0.6, n=12), with undetectable correlation. **(d)** Model of viscoelastic forces on k-fiber stubs due to microtubule (MT) crosslinking (magenta) and centromere (blue), and corresponding equation of motion (see Eq. (2) in text). **(e)** Solved equation of motion for relaxation of k-fiber post-ablation (top; see Eq. (3) in text) and cartoons of predictive contributions of model parameters (bottom). Steady state data showing a linear relationship between the steady state magnitude of relaxation (*A*) and stub length (*L*) (Fig. 1f) is consistent with model parameters predicting that the spindle elastic forces that are proportional to k-fiber length significantly contribute to force balance. Dynamic data that observed τ’s do not correlate with stub length (Fig. 3c), is consistent with model parameters predicting that viscous forces on k-fibers from their crosslinks to the spindle do not limit the timescale of relaxation, resulting in a timescale (*τ*) that is independent of *L*.

To project our observations of the post-ablation relaxation dynamics (Fig. 3c) and steady state (Fig. 1f) onto a framework for local anchorage mechanics, we built a simple phenomenological model and asked whether it was sufficient to recapitulate the main qualitative features of the data (Fig. 3d). This model provides a physical intuition for key factors that dominate the relaxation we observe (see Supplemental Experimental Procedures for details).

After ablation, the main spindle structural response is that of the centromere, along with the ablated k-fiber stub, relaxing toward the uncut sister k-fiber. The observation that long stubs can bear load (Fig. 1f) indicates that there must be crosslinkers between the k-fiber and spindle, since viscosity from the drag of the stub cannot alone bear steady state load. In turn, the non-zero relaxation time (Fig. 3c) indicates that at least one of the relaxing structures has a relevant viscous component. Thus, even the simplest model should account for material properties of both the centromeres and the k-fiber-to-spindle connections. To include both the elasticity and viscosity of the relaxing centromere, we model it as a Kelvin-Voigt solid [26]. We hypothesized that both elastic and viscous connections of the k-fiber to the spindle scale with k-fiber stub length. To model the frictional interaction of the k-fiber stub of known length with the spindle environment, we include a stub-length dependent viscous drag force. Finally, to account for transient binding crosslinkers between the k-fiber and the spindle environment, we include a stub-length dependent number of what we call “on/off bonds” that rupture at a particular force.

We assume no active response (e.g., spindle repair) during the short time scale observed. Pre-ablation, the system is in a stretched state, with the amount of stored stress indicated by the distance across the centromere. Post-ablation, the system relaxes with a time scale resulting from drag on the k-fiber stub and dissipation in the contracting chromosome. The change in distance across the centromere in response to ablation is equal to the deflection of the k-fiber stub, which we assume is stiff. Thus, matching the stresses (*σ*) on the centromere (left) and the k-fiber stub (right), we obtain:

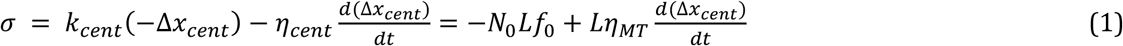

where *k*_*cent*_ is the centromere stiffness, *Δ*x*_*cent*_* is the extension of the centromere (beyond its rest length), *N_0_* is the density of k-fiber-to-spindle crosslinks per unit length, *L* is the k-fiber stub length, *f*_0_ is the maximum force generated per crosslink, *ηMT* accounts for viscous drag (per unit length) of the k-fiber stub in the spindle environment, and *η*_*cent*_ is the viscosity of the centromere. Simplifying,

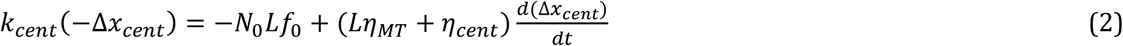

The solution to this equation is given by:

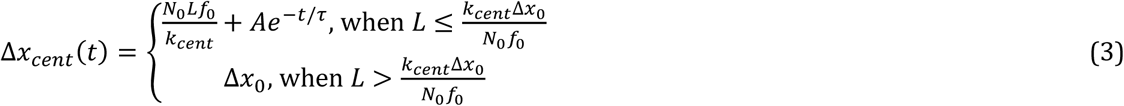

where

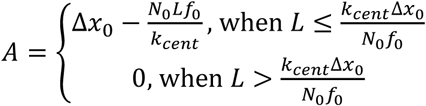

and

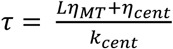

(Fig. 3e). Here, *A* describes the relaxation magnitude, 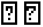 describes its timescale, and Δ*x*_0_ is the extension of the centromere at the time of ablation. Equation (3) qualitatively describes the main features of the dynamic and steady state relaxation we observe following ablation: the magnitude *A* scales linearly with k-fiber stub length *L* until it plateaus (Fig. 1f), and the relaxation follows single-exponential kinetics (Fig. 3b). The lack of correlation between the timescale of relaxation and stub length (Fig. 3c, e) suggests that the timescale of k-fiber movement in the spindle is dominated by centromere viscosity rather than viscous drag from the k-fiber-to-spindle connections, since the latter is expected to depend on stub length (Fig. 3e). If there is length-dependent viscosity (friction) between the k-fiber and rest of the spindle that significantly contributes to relaxation, we could not detect it amongst variation in measured *τ*’s, which may be a consequence of variation in material properties across centromeres. Together, the data suggest that k-fiber-to-spindle crosslinker bonds bear load in a k-fiber length-dependent manner, with longer microtubules more resistant to spindle deformation than short ones. Yet, the data suggest that k-fiber-to-spindle frictional connections do not limit the timescale for local k-fiber and spindle reorganization after acute spindle architectural changes. Thus, the local environment around a k-fiber is able to bear load yet fluid. The transient, dynamic nature of crosslinkers that would oppose a maximum force before falling off is compatible with this spindle property.

### NuMA-mediated microtubule crosslinking locally bears the load of chromosome movement in the spindle body

Non-k-fiber microtubules are a clear candidate structure to which k-fibers may anchor and share load: they are present [15] in the region where we map that load-bearing occurs (Fig. 1–2); they rapidly turnover [27], which would facilitate spindle reorganization; and they are mechanically connected to k-fibers [13]. We first tested whether increased microtubule-microtubule crosslinking led to more local load-bearing in the spindle. To do so, we increased crosslinking by treating cells with FCPT, which rigor binds the motor Eg5 to microtubules [28]. Eg5 localizes throughout the spindle [29], including in the vicinity where we observe local load-bearing, and functions as a microtubule slider and cross-linker [30]. We treated cells with concentrations of FCPT sufficient to decrease microtubule flux, indicating increased microtubule crosslinking, but that did not grossly perturb spindle structure or dynamics, allowing chromosomes to still move (Video 5; Fig. S1a; Supplemental Experimental Procedures). Under these conditions, we measured steady state centromere stretch after ablation near the kinetochore, where there is a larger dynamic range for measuring an increased ability to bear load following ablation. We observed decreased centromere relaxation following near-kinetochore ablation in FCPT (n=10, Fig. 4a, a’, Video 5), which demonstrates that increased microtubule-microtubule crosslinking is capable of increasing k-fiber anchorage and making load-bearing even more local.

**Figure 4.**
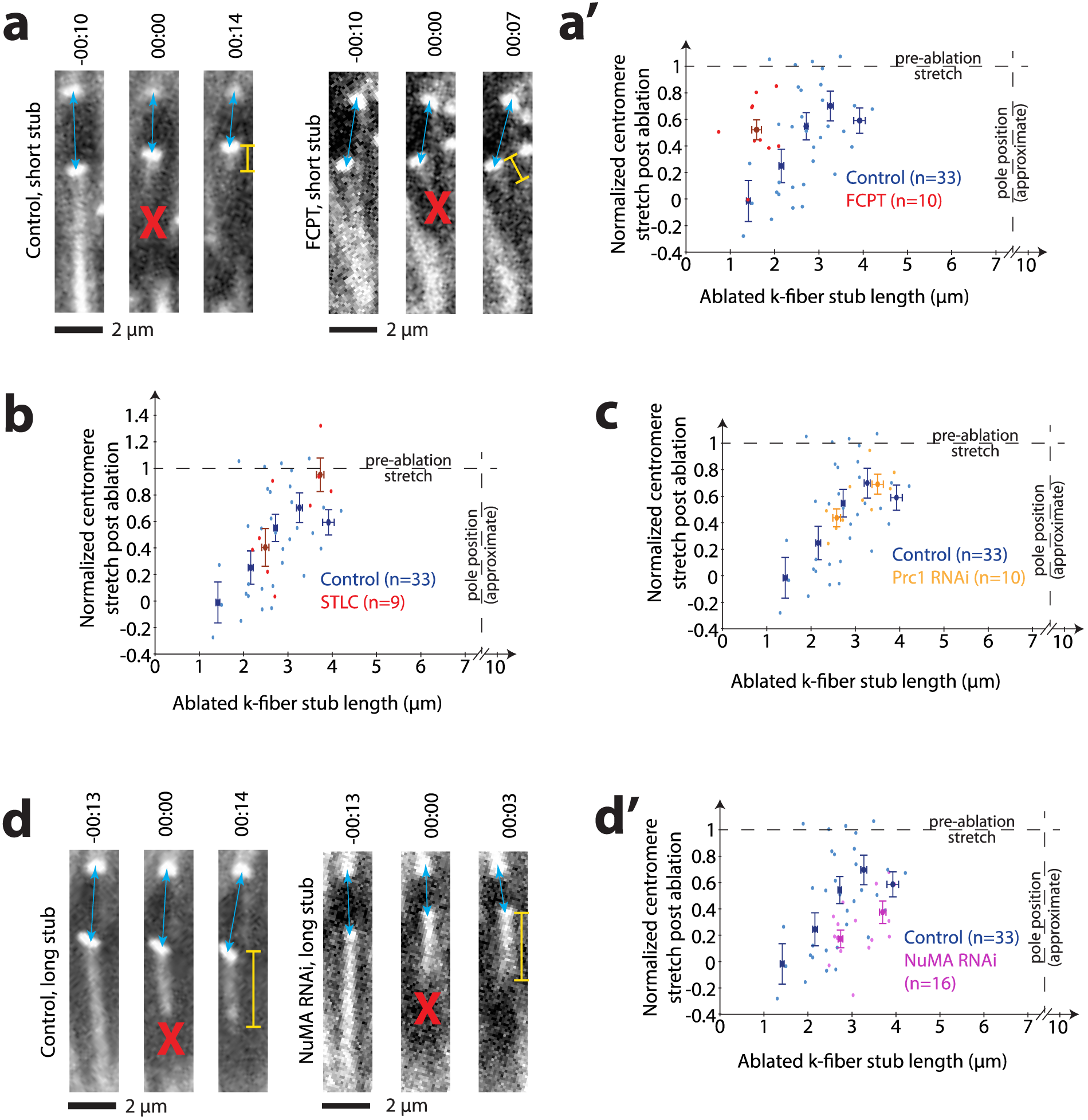
Local k-fiber anchorage can occur through microtubule crosslinking, and requires microtubule crosslinker NuMA but not PRC1 or Eg5. See also Fig. S1 and S2 and Videos 5-8. **(a)** Confocal live images of ablated (at red X) Ptk2 cells expressing EYFP-Cdc20 and EGFP-α-tubulin. Time since ablation, min:sec. Representative short k-fiber stubs (yellow bars) in a control cell (left, same movie as Fig. 1c) and in a cell treated with FCPT (right). Control centromere (blue arrows) relaxes, whereas in FCPT, centromere remains stretched. **(a’)** Normalized centromere stretch post-ablation vs. k-fiber stub length for control cells (blue) and cells treated with FCPT (red), which show increased load-bearing ability for short stubs. **(b)** Normalized centromere stretch post-ablation vs. k-fiber stub length for control cells (blue) and cells treated with STLC (red), which show similar load-bearing ability. **(c)** Normalized centromere stretch post-ablation vs. k-fiber stub length for control cells (blue) and cells depleted of PRC1 by RNAi (gold), which show similar load-bearing ability. **(d)** Imaging as in (a). Representative long k-fiber stubs in control cell (left, same movie as in Fig. 1d) and in a cell depleted of NuMA by RNAi. Control centromere (blue arrows) remains stretched, whereas in NuMA depletion, centromere relaxes. **(d’)** Normalized centromere stretch post-ablation vs. k-fiber stub length for control cells (blue) and cells depleted of NuMA by RNAi (magenta), which show a decreased load-bearing ability for long stubs. In all graphs, light dots, individual measurements; dark dots, binned data; error bars, s.e.m.

Given that microtubule crosslinking is able to locally anchor k-fibers, we next asked if key spindle microtubule crosslinkers played essential roles in locally bearing load along k-fibers. This time, we focused on ablation far from the kinetochore, where the significant load-bearing by a longer stub gave the greatest dynamic range for detecting a decreased ability to bear load. We first examined the role of the active force generator Eg5, which, in addition to its crosslinking role, exerts outward force in the spindle [31], and has been previously reported to mechanically connect neighboring chromosome movements [18]. When we examined bipolar spindles treated with STLC [32], which causes Eg5 to release from microtubules, we saw no effect on the k-fiber’s load-bearing ability (n=9, Fig. 4b, Video 6, Fig. S1b). Thus, while Eg5 is capable of increasing the density at which load-bearing force is generated when its microtubule affinity is increased, it is not essential for local load-bearing in the spindle body.

Second, we examined the role of the passive microtubule crosslinker PRC1, which bundles microtubules in the center of the spindle where local load-bearing occurs. It has a reported role specifically in “bridging fibers” [5] whose geometry – connecting sister k-fibers by spanning the centromere [33] – is well-suited to maintaining poleward force on chromosomes [15, 34]. Furthermore, overexpression of PRC1 and concomitant thickening of bridging fibers results in a greater angular deflection of the centromere following ablation in HeLa cells [5]. However, following depletion of PRC1 by RNAi [35], we did not detect any effect on local k-fiber load-bearing in Ptk2 cells (n=10, Fig. 4c, Video 7, Fig. S1c), indicating that PRC1 is not essential for local load-bearing. Thus, it may be that PRC1-mediated bridging microtubules can effectively stiffen the centromere but do not significantly bear the load of stretching it, or there may be species-specific differences.

Finally, we examined a role for NuMA, a microtubule crosslinker that stabilizes spindle poles [3], where it primarily localizes, but that also localizes diffusely throughout the spindle body, including on k-fibers [36] (Fig. S2). Since complete NuMA depletion significantly perturbs spindle structure, making it difficult to differentiate between a direct or indirect role of NuMA in load-bearing, we focused on cells that exhibited only mild phenotypes of NuMA knockdown, such as mildly splayed poles or slightly detached centromeres (Video 8, Fig. S1d). Strikingly, even these cells displayed a decrease in local load-bearing for long k-fiber stubs (n=16, Fig. 4d, d’, Video 8, Fig. S1d) indicating a specific role for NuMA in local load-bearing and k-fiber anchorage. This partial rather than complete decrease in load-bearing may stem from residual NuMA, as expected from our cell selection protocol, or from possible redundant anchoring molecules. NuMA’s role may either be through its own passive microtubule cross-linking ability, or through its role in helping the minus-end directed motor dynein attach to microtubules as cargo [37]. The observation that the load-bearing map does not change without PRC1 (Fig. 4c) suggests that NuMA’s role is not specific to PRC1-mediated bridging fibers.

Our work raises the question of why NuMA can locally bear the load of chromosome movement – so that its absence changes the spindle’s load-bearing map – while other microtubule crosslinkers Eg5 and PRC1 do not. We speculate about two possible mechanical advantages for NuMA for this function. First, Eg5 and PRC1 and/or their homologs have a preference for linking antiparallel microtubules [38–41], while we are unaware of any such known preference for NuMA, and indeed, NuMA’s role in astral arrays implies the capability of linking parallel microtubules. Parallel microtubule crosslinking, which may thus be more efficiently mediated by NuMA than by Eg5 and PRC1, may be preferred for k-fiber anchorage. Second, interestingly, NuMA exerts more friction under load directed toward the plus-end of microtubules (as would be expected in opposition to force at kinetochores) than toward the minus-end [42]; such a preference does not exist for PRC1 [42], and is, to our knowledge, unknown for Eg5. However, whether NuMA’s frictional asymmetry is key to its function is not yet clear.

Altogether, the data herein suggest a model where NuMA distributes the load of chromosome movement from k-fiber microtubules to nearby spindle microtubules through NuMA-based microtubule-microtubule crosslinking (Fig. 5). While NuMA’s primary known functions have been at poles and at the cell cortex [43], our work suggests a function for NuMA within the spindle body. We propose that NuMA contributes to dense crosslinking of microtubules throughout the spindle, locally bearing load both around chromosomes and at poles, and mechanically isolating these from each other.

**Figure 5.**
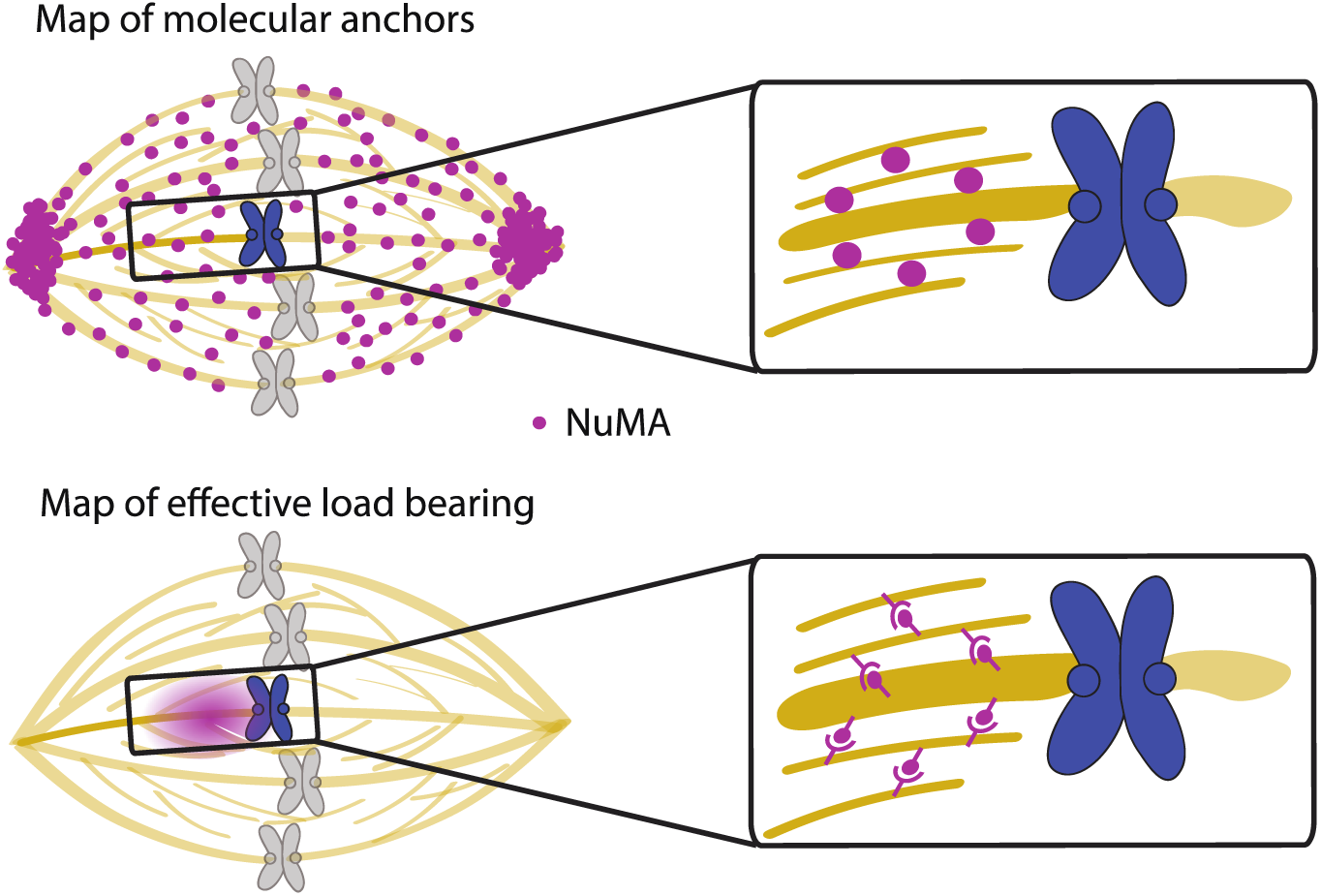
Model of k-fiber load-bearing. The data herein suggest that widespread NuMA-mediated crosslinks in the spindle (top, purple circles) provides local load-bearing (bottom, purple cloud). Key to this model, NuMA acts in all spindle regions, locally bearing load everywhere and providing mechanical isolation and redundancy that are well-suited to support robust chromosome segregation.

### Local k-fiber load-bearing ensures mechanical redundancy and robust function

The proportionality of elastic force to k-fiber stub length, and the smooth, exponential relaxation we observe over time after ablation, suggest a dense network of anchoring microtubules: indeed, in a sparse network we would instead expect discrete relaxation events as individual crosslinks ruptured. Unlike a sparser network, where each connection would be mechanically crucial, a dense network provides many local “backup” paths of force transmission for bearing the load of chromosome movement. In this way, neither connections of chromosomes to the spindle nor global spindle architecture would be jeopardized by events, such as chromosome movements, that require dynamic turnover of local connections of k-fibers to the spindle. Indeed, chromosomes must move past each other as they move to and around the metaphase plate, and the spindle must dramatically reorganize itself as it progresses through mitosis. Notably, the ability of the spindle to locally bear load implies that force generation on kinetochores, thought to for example stabilize kinetochore-microtubule attachments [44], is robust to distant changes in spindle architecture or forces. Thus, local and redundant load-bearing could allow the mammalian spindle to dynamically reorganize itself while preserving its ability to bear the load of chromosome movement, and is well-suited to support chromosome movement in species where few microtubules directly connect the kinetochore and spindle pole [45]. Looking forward, it will be important to map whether and how load-bearing is regulated amongst different microtubule populations (e.g., k-fiber and interpolar microtubules), and whether the spindle holds on to k-fiber microtubules specifically. It will also be of interest to test how load is distributed in space and time across spindles from different species, and how this relates to the functions – and diverse or similar mechanical solutions – of these spindles.

## Experimental Procedures

### Cell culture, transfection, siRNA, and small molecule treatment

See Supplemental Experimental Procedures for details. Briefly, WT Ptk2 cells and Ptk2 GFP-α-tubulin cells (gift of A. Khodjakov, [46]) were cultured in MEM and imaged in phenol free MEM (Invitrogen) as previously described [16]. For imaging of kinetochores and microtubules, Ptk2 cells stably expressing EYFP-Cdc20 (gift of J. Shah) were transfected with GFP-α-tubulin using Fugene 6 (Promega). For depletion of NuMA or PRC1, cells were also transfected with siRNA for NuMA or PRC1 using Oligofectamine (Life Technologies, Carlsbad, CA) as previously described [35]. For speckle imaging of microtubules, Ptk2 cells were simultaneously transfected with K560rig-SunTag24x-GFP and scFv-GCN4-GFP [24] using Fugene 6.

To increase anchorage by rigor binding Eg5, we treated with 20 μM FCPT (gift of T. Mitchison) for 15-30 min. Slowing of microtubule flux verified drug effectiveness. To inhibit Eg5 motor activity, we treated with 5 μM S-trityl-L-cysteine (STLC, Sigma) in MEM for 30 min. The presence of monopolar spindles verified drug effectiveness. For speckle experiments, we used SiR tubulin to simultaneously visualize microtubules. We treated with 10 μM verapamil and 100 nM SIR-tubulin (Spirochrome) [47] for 2-6 h to allow time for incorporation of SIR-tubulin into spindles. Under these conditions, there was no detected defect in spindle appearance or behavior.

### Imaging and targeted laser ablation

See Supplemental Experimental Procedures for details. Briefly, live imaging was performed on a Nikon Eclipse Ti-E inverted microscope with a Yokogawa CSU-X1 spinning disk confocal, 405/488/561/642 nm diode lasers, Chroma and Semrock filters and an Andor iXon3 camera. Cells were imaged by phase contrast and fluorescence with a 100X 1.45 Ph3 oil objective through a 1.5X lens. Cells were imaged at 29-31°C and 5% CO2 in a Tokai Hit scope top incubator, using the Nikon Perfect Focus System. Targeted laser ablation using 551 nm light was performed using a galvo-controlled MicroPoint Laser System (Photonic Instruments) operated through Metamorph. We verified ablation through depolymerization of microtubule plus-ends created by k-fiber severance, or through the ensuing relaxation response when it was present.

### Data analysis

See Supplemental Experimental Procedures for details. Briefly, k-fiber ends, spindle pole positions, microtubule speckles, and photobleached spots (for quantifying flux) were tracked manually in home-written Matlab (R2012a/R2016a) programs. All analysis based on these positions and plots were created with home written Matlab programs. Kymograph of fast relaxation (Fig. 3) was created in ImageJ [48].

### Modeling

Equation of motion for viscoelastic relaxation (Fig. 3d) was solved by hand analytically. For derivation, see Supplemental Experimental Procedures.

## Author contributions

M.W.E. and S.D. designed experiments. M.W.E. and S.D. obtained preliminary data; D.U. designed and validated the strategy for RNAi knockdown of NuMA and PRC1 in Ptk2 cells; M.W.E. performed all other experiments reported here, and all analyses. M.W.E. and M.P. designed and performed modeling. M.W.E. and S.D. wrote the paper.

## Acknowledgments

We thank Alexey Khodjakov for Ptk2 GFP-α-tubulin cells, Jagesh Shah for Ptk2 EYFP-Cdc20 cells, Marvin Tanenbaum and Ron Vale for K560-SunTag plasmids, Tim Mitchison for FCPT, and Tim Mitchison and the HMS NIC who provided support to obtain preliminary data. We thank Ron Vale, Dan Needleman, Tim Mitchison, Megan Valentine, and Christina Hueschen, Jon Kuhn and other members of the Dumont lab for helpful discussions.

This work was supported in part by the National Institutes of Health (R00GM09433, DP2GM119177). S.D. is a Rita Allen Scholar and Searle Scholar. M.P. is supported by the W.M. Keck Foundation. M.W.E. is a Damon Runyon Fellow supported by the Damon Runyon Cancer Research Foundation (DRG-2170-13).

The authors declare no competing financial interests.

